# Genetic diversity analysis of an isolate of microsporidia infesting *Athetis lepigone*

**DOI:** 10.1101/526368

**Authors:** Haijian Zhang, Jian Song, Yunhe Yang, Jingjie An, Hongxia Ma, Yaofa Li

## Abstract

In this study, PCR amplification, cloning and sequencing analysis were adopted to explore genetic diversity of microsporidia (LEP9557) infecting *Athetis lepigone*. The small subunit ribosomal DNA (SSU rDNA), internal transcribed spacer (ITS) and intergenic spacers (IGS) of ribosomal RNA (rRNA) were cloned from the strain and sequenced. By means of multiple sequence alignment, we found that the three gene regions had different levels of polymorphism. There was greater polymorphism in ITS (74 variable sites) and IGS (55.59%) regions than in the SSU rDNA (17 variable sites). Phylogenetic analysis was performed using Kimura 2-parameter with neighbor joining and the results showed that LEP9557 had a close relationship with *Nosema bombycis*. Sequences of each clone were submitted to Genbank (Accession number: MF150254-MF150258). All of the results indicated the presence of genetic diversity in LEP9557, which established the foundation for identifying the phylogeny and relationships with other microsporidian strains, and had significant biological meaning for maintaining the survival and population continuity of the strain.

---

*Athetis lepigone* Möschler (Lepidoptera: Noctuoidea) is a world-wide distributed noctuid insect. In the past decade, *A. lepigone* broke out in Northern China and caused significant losses in summer maize [1,2]. Chemical pesticides are the conventional method used against this pest. However, excessive pesticides would cause various environmental problems. Therefore, it is necessary to develop environmentally-acceptable alternative approaches to control this pest in corn fields [3].

Microsporidia, a group of obligate intracellular worldwide distribute parasitic fungi, which can infect a variety of hosts including vertebrates and invertebrates [4]. As a common pathogen of economic insects and animals, some species of microsporidia take a heavy economic toll on the agricultural industry, such as sericulture. However, microsporidia can also be used as efficient biological control agents against several agricultural pests. Ran et al. [5] described a previously unknown microsporidian pathogen (*Nosema sp*.), which can invaded *Helicoverpa armigera* Hübner (Lepidoptera: Noctuidae) larvae and could be transmitted to their offspring, causing great morphological changes in *H. armigera* tissues, such as midgut, fat bodies, malpighian tubules and nerve cells. Moreover, from another tests carried out at Montana, USA showed that, high level of *Nosema locustae* Canning (Dissociodihaplophasida: Nosematidae) caused significant reduction in grasshopper densities during the season of treatment and the two following subsequent seasons when applied to rangeland by aircraft at the dosage of 2.1 × 10^8^ spores per ha [6]. As a successful biocontrol agent, *Nosema* has been extensively studied and widely used because of its unique transmission pathways, low consumption, persistence and environmentally friendly property.

In our previous study, a microsporidian strain has been isolated from overwintering larvae of *A. Lepigone* [4,7], which had the closest relationship with *Nosema bombycis* and was considered to be potentially applied to control *A. lepigone* because of its high infectivity in larvae of *A. lepigone*. After being infected, *A. lepigone* larvae did not die directly, but showed series of symptoms such as anorexia, growth restriction, incomplete molting, pupation deformity, eclosion deformity and decreasing fecundity. Thus, this microsporidia strain is worthy of further study in sustainable *A. lepigone* control.

Ribosomal DNA is usually used for examining relationships among all known life forms, because it contains highly variable regions including many of the nucleotide positions in internal transcribed spacer (ITS), intergenic spacers (IGS) and highly conserved regions such as portions of the SSU rDNA and LSU rDNA. SSU rDNA is often used for the analysis of relationships between genera, and therefore can be used to determine the taxonomic status of microsporidia at generic level. Having a greater variation in biological species and subspecies, rDNA ITS could be used for the species level study [8].

In this study, we selected an isolate of *Nosema* (LEP9557) infecting *A. lepigone* and obtained its rDNA sequences including SSU, ITS and IGS regions to evaluate its genetic diversity. We also isolated and sequenced multiple clones (from 5 to 16) for each of the three rDNA regions from this population. Considering the variations in the ITS and IGS regions, this information might be very useful for the research in continuation and evolution of microsporidian species and for the biocontrol of *A. lepigone* by microsporidia.

## 1 Materials and methods

### 1.1 Spore isolation and purification

To ensure the same source of infection, we obtained the microsporidian spores from an infected *A. lepigone* larva collected from Handan, Hebei province, China. The third-instar larvae of *A. lepigone* in captivity were fed on the spores at a concentration of 1.0×10^7^ spores/mL for 40 h, and then reared on artificial diet for 6-7 d. Subsequently, the midguts of infected larvae were separated, filtered and centrifuged to obtain the crude extracts of the microsporidia. Spores were purified by a discontinuous Percoll density gradient (25%, 50%, 75%, 100%, v/v), centrifuged at 17,500 rpm for 20 min, rinsed several times, and stored at 4°C [9].

### 1.2 Amplification, cloning and sequencing of rDNA sequence

Genomic DNA was extracted using CTAB kit (Aidlab Biotechnologies Co., Ltd Beijing China). Three pairs of primers corresponding to ITS, SSU, IGS rDNA (Table 1) were designed and synthesized according to the previous report [10]. The composition of the PCR reactions: 12.5 μL 2 × EasyTaq Supermix; 0.5μL each of the forward and reverse primers (10 μM); 1 μL template DNA; 10.5 μL ddH_2_O. The amplification was carried out using the following conditions: an initial DNA denaturation step at 94°C for 5min followed by 30 cycles of denaturation at 94°C for 1 min; annealing for 30 s (SSU at 53°C, ITS at 58°C, IGS at 50°C), and elongation at 72°C for 45 s. The last step was a 8 min extension at 72°C. Then the PCR products were purified using the Agarose Gel Extraction kit (TIANGEN, Beijing, China), and transformed into *Escherichia coli* DH5α. Three replicates of each clone were sent to the Sangon Biotech Company for sequencing.

**Table 1.**
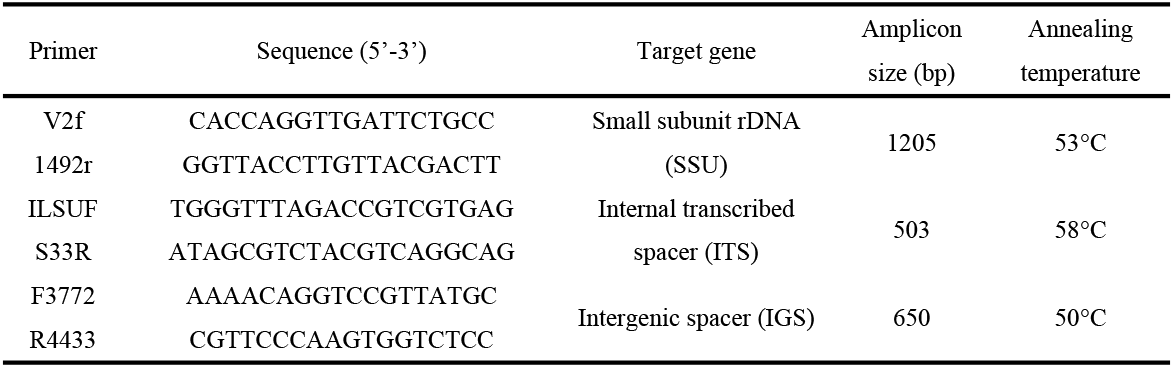
Primers used for amplification and sequencing of rDNA

### 1.3 Sequence analysis

Sequences obtained from each clone were identified by BLAST of the NCBI GenBank. Phylogenetic trees of SSU, ITS and IGS rDNA were constructed using the neighbor-joining method (Saitou and Nei, 1987) in MEGA 5.2 program. Bootstrap support was evaluated based on 1,000 replicates.

## 2 Results

### 2.1 SSU rDNA sequences

We obtained five clones of SSU rDNA (S1-S5). The length of SSU rDNA clones had little variations, except S4 which was 1,209 bp, the other four clones were all 1,208 bp. The sequences of the five clones were deposited in GenBank, with the GenBank accession numbers of MF150254-MF150258 (S1-S5) (Table 2). The sequence similarity was 99% among these five clones. There were only 17 polymorphic sites including 16 transitions / transversions and one deletion at position 738 (Table 3).

**Table 2.** The sequences lengths of five clones of SSU rDNA

| Clones<sup>a</sup> | Length (bp) | GenBank Accession Number |
| --- | --- | --- |
| S1 | 1208 | MF150254 |
| S2 | 1208 | MF150255 |
| S3 | 1208 | MF150256 |
| S4 | 1209 | MF150257 |
| S5 | 1208 | MF150258 |

<sup>a</sup> S1-S5 indicated respectively small subunit rDNA for isolate LEP9557

**Table 3.** Seventeen polymorphic sites in the SSU rDNA among LEP9557

| Clones<sup>a</sup> | Positions of varied nucleotide |  |  |  |  |  |  |  |  |  |  |  |  |  |  |  |  |  |  |
| --- | --- | --- | --- | --- | --- | --- | --- | --- | --- | --- | --- | --- | --- | --- | --- | --- | --- | --- | --- |
|  | 1<sup>b</sup> | 95 | 160 | 297 | 369 | 388 | 543 | 605 | 637 | 649 | 738 | 793 | 960 | 985 | 1004 | 1085 | 1094 | 1107 | 1209<sup>b</sup> |
| S1 | T | C | G | T | A | T | A | T | A | C | -<sup>c</sup> | G | G | T | A | T | T | C | G |
| S2 | T | T | G | T | A | C | A | T | A | T | - | A | G | T | A | T | T | C | G |
| S3 | T | C | A | A | A | T | A | C | A | T | - | A | G | A | T | C | T | C | G |
| S4 | T | C | A | A | G | T | G | T | A | T | - | A | G | T | A | T | T | T | G |
| S5 | T | C | A | A | A | T | A | T | G | T | A | A | A | T | A | T | C | T | G |

<sup>a</sup> S1-S5 indicated respectively small subunit rDNA for isolate LEP9557
<sup>b</sup> indicated start and end sites, respectively.
<sup>c</sup> “-” indicated a missing nucleotide

Phylogenetic analysis (Fig 1) showed that these five clones clustered separately and then gathered together to form a subgroup which shared 59% similarity with AB097401.2 *Nosema bombycis* but had a relatively distant phylogenetic relationship with FJ772435.1 *Nosema heliothidis* and AY747307.1 *Nosema spodopterae*. This result preliminarily indicated that LEP9557 belonged to *Nosema* genus.

**Fig 1.**
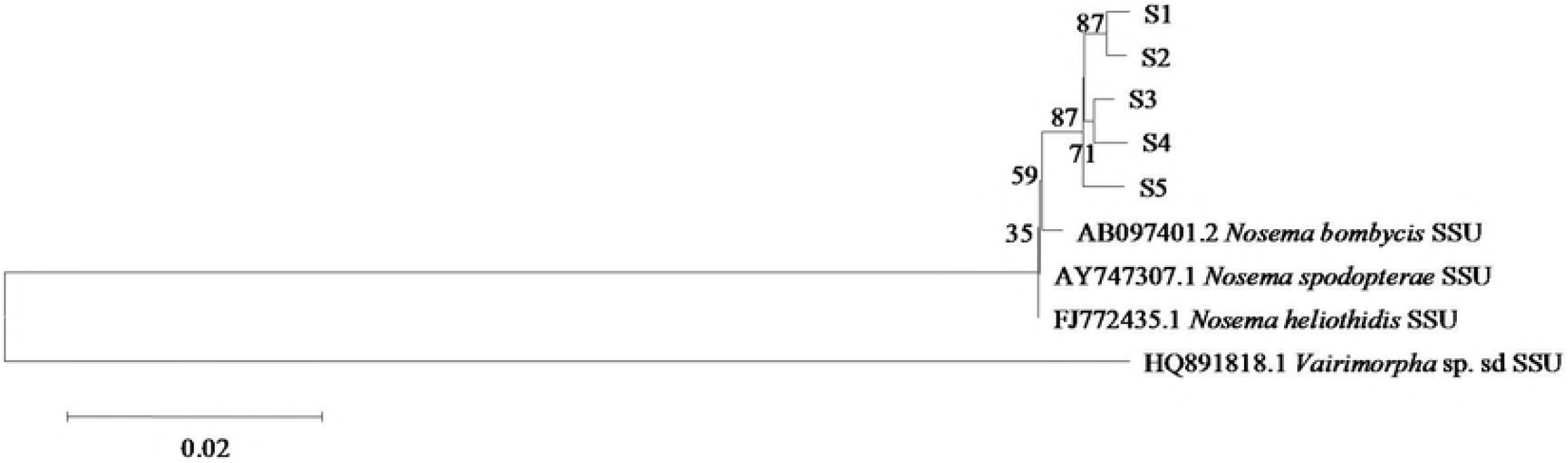
Phylogenetic analysis of isolate LEP9557 based on SSU rDNA sequences.

### 2.2 ITS rDNA sequences

The sequence lengths of the sixteen clones of rDNA ITS ranged from 180 to 188 bp (Table 4), suggesting obvious length polymorphism. Polymorphism analysis revealed that, each sequence of the sixteen clones was unique with 96%~99% similarity to *N. bombycis* rDNA ITS sequence from GenBank. Sequence alignment showed that conversions / transversions and insertions / deletions existed on mononucleotide of ITS gene. There were 74 variation sites including 25 insertion/deletion sites and most of the variation sites were within the ITS sequence (Fig 2).

**Fig 2.**
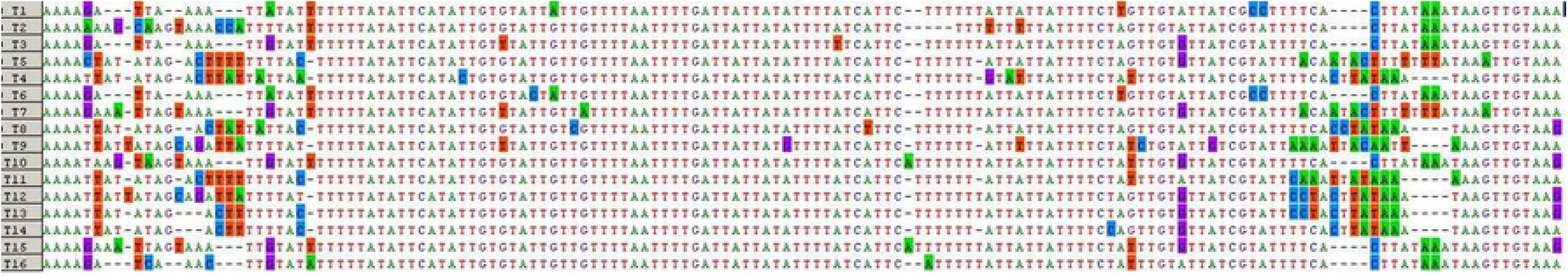
Polymorphism analysis of rDNA ITS of LEP9557. Colored regions indicate the difference of the sixteen individuals.

**Table 4.** The sequences lengths of sixteen clones of rDNA ITS

| Clones<sup>a</sup> | Length(bp) | GenBank Accession Number |
| --- | --- | --- |
| T1 | 180 | MF150259 |
| T2 | 180 | MF150260 |
| T3 | 180 | MF150261 |
| T4 | 180 | MF150262 |
| T5 | 183 | MF150263 |
| T6 | 183 | MF150264 |
| T7 | 184 | MF150265 |
| T8 | 184 | MF150266 |
| T9 | 184 | MF150267 |
| T10 | 185 | MF150268 |
| T11 | 185 | MF150269 |
| T12 | 186 | MF150270 |
| T13 | 186 | MF150271 |
| T14 | 187 | MF150272 |
| T15 | 188 | MF150273 |
| T16 | 188 | MF150274 |

<sup>a</sup> T1-T16 indicated respectively rDNA ITS for isolate LEP9557.

We performed phylogenetic analysis of the ITS sequence of LEP9557 shown above (Fig 3). T1 and T6 were in the same branch with 90% similarity and they had close relationship with JF443603.1 *N. bombycis*. Similarly, T16, T5 and T9 also had high similarity with *N. bombycis*. In addition, it can be observed from the phylogenetic tree that microsporidian species in the *Nosema* genus had obvious genetic diversity (Fig 3). For example, GQ334400.1 *N. bombycis* and FJ443628.1 *N. heliothidis* were in the same branch with only 91% similarity concluding that, LEP9557 also had a certain degree of genetic diversity because the similarities between any two ITS clones were low. This phenomenon may be due to the short size and high mutation rate of ITS sequence.

**Fig 3.**
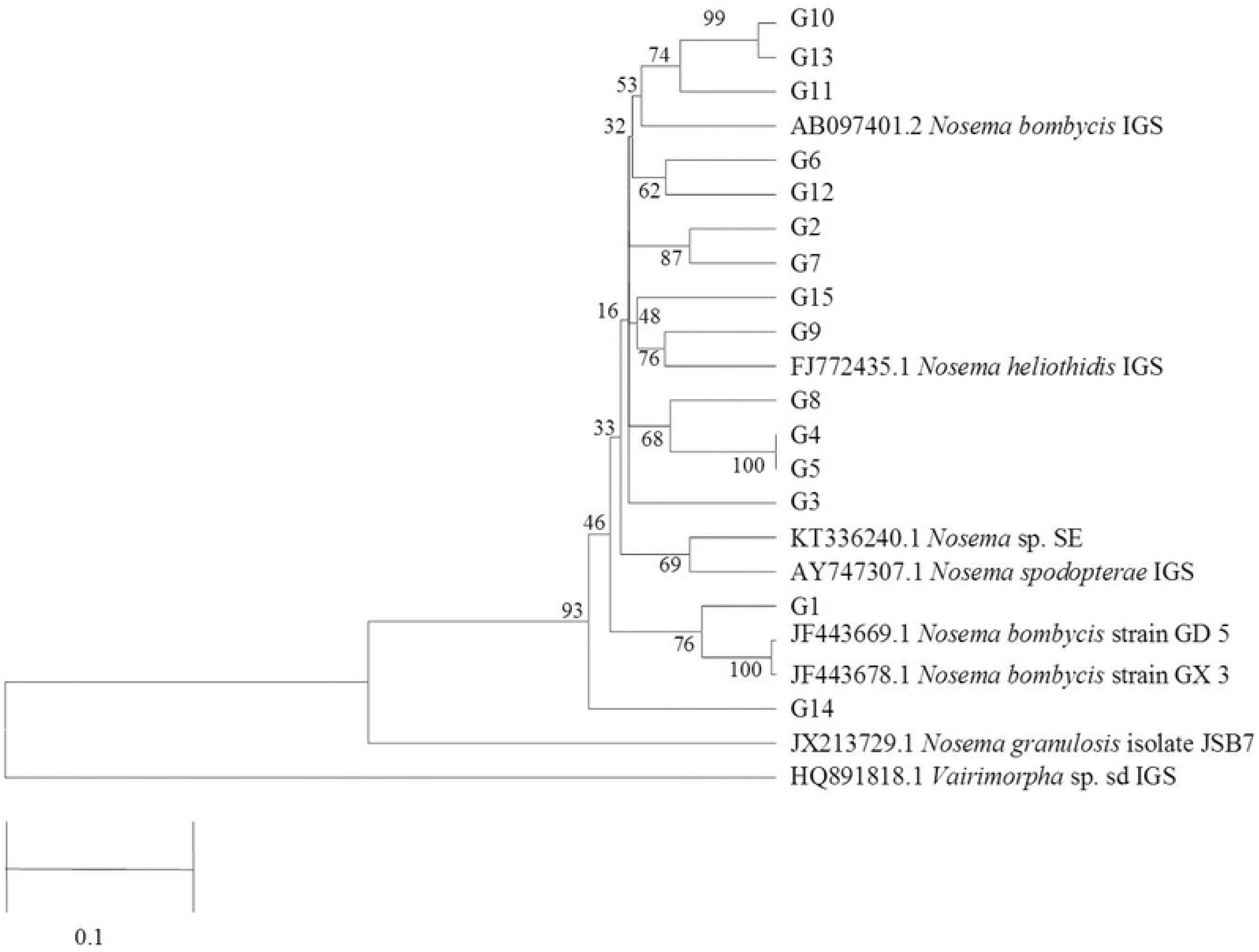
Phylogenetic analysis of ITS rDNA.

### 2.3 Analysis of IGS rDNA sequences

The fifteen IGS sequences of the isolate varied in length from 269 bp to 286 bp (Table 5), showing obvious length polymorphism, with about 55.59% polymorphic positions. Among these clones, G4 and G5 had same sequence and their sequence similarity with *N. bombycis* rDNA sequence from GenBank was between 93%~99%. Sequence alignment showed that the IGS region was highly variable (Fig 4). Base sites were changeable including conversions/transversions of mononucleotides and insertions/deletions of bases, and most of the polymorphic positions were located within the IGS sequences.

**Fig 4.**
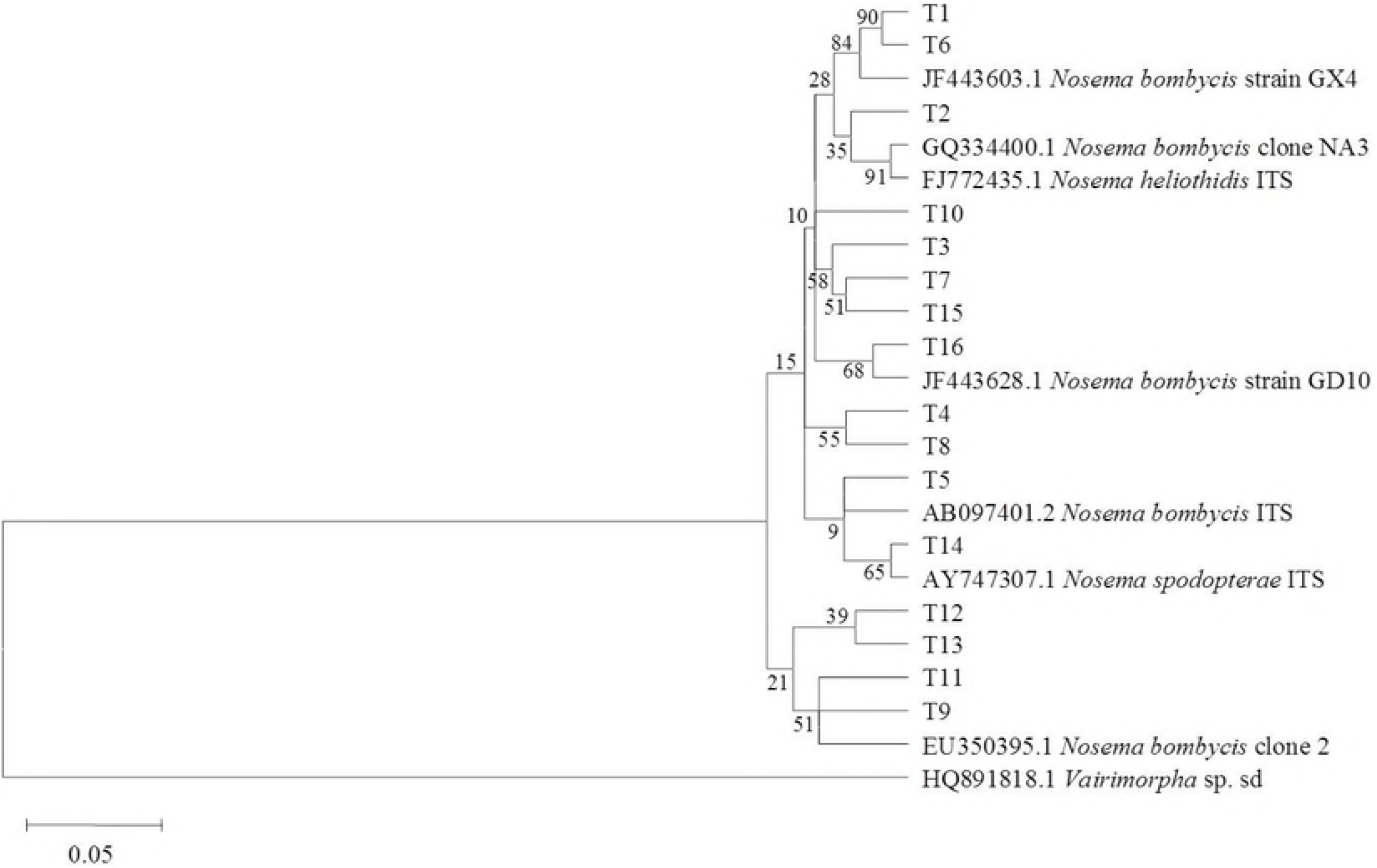
Polymorphism analysis of rDNA IGS of LEP9557. Colored regions indicate the difference of the sixteen individuals.

**Table 5.** The sequences lengths of fifteen clones of rDNA IGS.

| Clones<sup>a</sup> | Length(bp) | GenBank Accession Number |
| --- | --- | --- |
| G1 | 275 | MF150275 |
| G2 | 279 | MF150276 |
| G3 | 278 | MF150277 |
| G4 | 281 | MF150278 |
| G5 | 281 | MF150279 |
| G6 | 276 | MF150280 |
| G7 | 278 | MF150281 |
| G8 | 283 | MF150282 |
| G9 | 275 | MF150283 |
| G10 | 283 | MF150284 |
| G11 | 280 | MF150285 |
| G12 | 269 | MF150286 |
| G13 | 280 | MF150287 |
| G14 | 281 | MF150288 |
| G15 | 286 | MF150289 |

<sup>a</sup> G1-G15 indicated respectively rDNA IGS for isolate LEP9557

Based on the phylogenetic tree of LEP9557 IGS rDNA, the similarities between G10 and G13 and between G4 and G5 were up to 99% and 100%, respectively (Fig 5). This result together with the result described above indicated that, G4 and G5 were the copy of same gene. This result was similar in both the IGS and the ITS region, which, again, indicated that this isolate had intraspecific genetic diversity and its IGS region also had high mutation rate.

**Fig 5.**
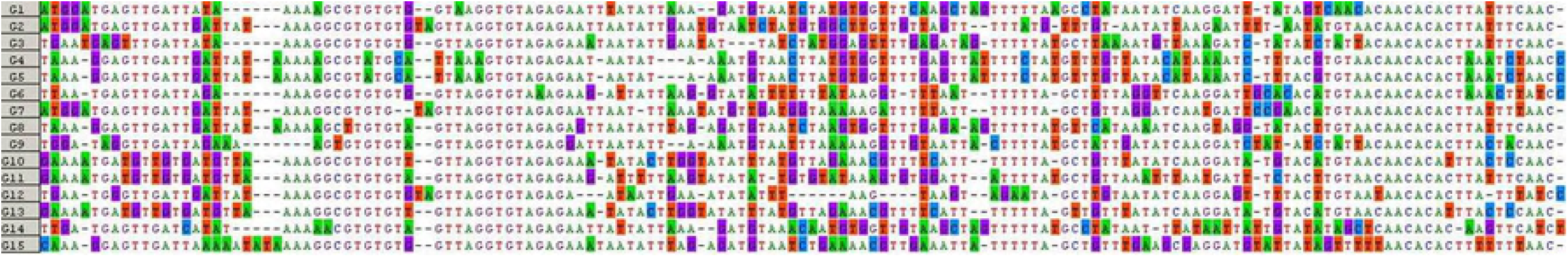
Phylogenetic analysis of IGS rDNA.

## 3 Discussion

At present, most of the studies on molecular biology of microsporida are focused on rDNA genes that are often present as multiple copies of tandem repeat units in all of the microsporidian genomes [11]. Each of the repeat units of ribosomal genes was composed of transcription units and intergenic spacer regions. For LEP9557, the pattern of the transcript unit was LSU-ITS-SSU-IGS-5S. This arrangement is similar to that of ribosomal genes of *N. bombycis* and *N. antheraea* [12].

Cunning et al. [13] indicated that the SSU rDNA and other highly conserved genes may be useful in the classification between genera rather than between species. Tsai et al. [14] suggested that ITS rDNA should have great potential applications in the classification between closely related species. The result of our study revealed that, there was only 1 bp difference between the sequences of the SSU rDNA gene. Polymorphism of mononucleotides mainly existed among SSU rDNA clones, while the mutation frequencies of regions such as rDNA-ITS and rDNA-IGS were higher than that of SSU rDNA. This might because SSU rDNA is extremely conservative and restricted when the close systematic relationship was measuring.

ITS sequences have more differential positions owing to a faster evolutionary rate. This polymorphism could be used for fine classification of species and subspecies and as a specific-fingerprinting to study variations in species [15]. The high variability of ITS has unique advantage in the study of biological classification, identification, phylogenetic and structural changes, and has been widely used in the field of clinical medicine [16], applied microbiology [17,18] and ecology [19,20]. In this study, ITS sequences, with high variability, were different between these clones not only in length but also in sequence. The difference in length and sequence of ITS would play an important role in the identification of PCR products. Considering that, if the ITS sequences were less than 300 bp, the information we obtained from those sequences were limited. With short sequences, different clones were easily clustered in phylogenetic analysis. If full-length sequences of rDNA ITS and IGS were used in the genetic polymorphic analysis of microsporidia, the possibility of deviation could be reduced.

In this study, LEP9557 represent significant intraspecific genetic differentiation, which may be a result of genetic variation during passage or the specificity of host tissues in which population individual parasite dwells. Sequence polymorphism showed the diversity and complexity of the genetic background of the microsporidian isolate. This result may be an adaptive reaction to resist adverse environment during evolution, and may have biological significance for maintaining the survival and continuity of the microsporidian population.

## Acknowledgments

The authors are grateful to Tao Zhang, Key Laboratory of IPM on Crops in Northern Region of North China, Ministry of Agriculture, for assistance in revising this article.

## References

1. Shan XN, Yang PY, Zhao ZH, Yan DL, Wang XY. Causes and prevention measures of *Athetis lepigone* in corn field during 2011. Chin Plant Prot. 2011; 31 (8): 20–22.

2. Wang ZY, Shi J, Dong JG. Reason analysis on Proxenus lepigone outbreak of summer corn region in the Yellow River, Huai and Hai Rivers Plain and the counter measures suggested. J Maize Sci. 2012; 20(1): 132–34.

3. Wasson K, Peper RL. Mammalian microsporidiosis. Vet Pathol. 2000; 37(2): 113–28.

4. Zhang HJ, Song J, Du LX, Yang YH, Shi J. Control Efficacy of One Microsporidian against *Athetis lepigone* Larvae. Chin J Biolo Control. 2016; 32(4): 462–67.

5. Ran HF, Feng SL, Pan WL, Fan XH. Observations on histopathology of tissue infected by Nosema sp. (Microsporida) in *Helicoverpa armigera* (Hǜbner) larvae. Acta Entomol Sinica. 2003; 46(1): 118–21.

6. Henny JE, Onsager JA, Large-scale test of control of grasshoppers on rangeland with Nosema locustae. J Econ Entomol. 1982; 75(75): 31–35.

7. Yang YH, Shi J, Zhang HJ, Guo N, Du LX. Pathogenic Mechanism of a Microsporidium Strain Isolated from *Athetis lepigone*.Chin J Biolo Control. 2017; 33(4): 571–74.

8. Niu QL, Luo JX, Yin H. Advances and application of rDNA-ITS in the molecular taxonomy of parasite. Chin J Vet Parasitol. 2008; 16(4), 41–47.

9. Gatehouse HS, Malone LA. The Ribosomal RNA Gene Region of *Nosema apis*(Microspora): DNA Sequence for Small and Large Subunit rRNA Genes and Evidence of a Large Tandem Repeat Unit Size. J Invertebr Pathol. 1998; 71(2): 97–05.

10. Huang WF, Tsai SJ, Lo CF, Soichi Y, Wang CH. The novel organization and complete sequence of the ribosomal RNA gene of *Nosema bombycis*. Fungal Genet Biol. 2004; 41(5): 473–81.

11. Brugere JF, Cornillot E, Metenier G, Bensimon A, Vivares CP. Encephalitozoon cuniculi (Microspora) genome: physical map and evidence for telomere-associated rDNA units on all chromosomes. Nucleic Acids Res, 2000; 28(10): 2026–33.

12. Wang L, Chen K, Yao Q, Gao G, Zhao Y. The sequence and phylogenetic analysis of the ribosomal RNA gene and the ITS region of *Nosema antheraeae*. Sci Agric Sin. 2006; 39(8): 1674–79.

13. Canning EU, Curry A, Cheney S, Lafranchi-Tristem NJ, Haque MA. Vairimorpha *imperfect* n. sp., a microsporidian exhibiting an abortive octosporous sporogony in Plutella *xylostella* L. (Lepidoptera: Yponomeutidae). Parasitology. 1999; 119(3): 273–86.

14. Tsai SJ, Lo CF, Soichi Y, Wang CH. The characterization of Microsporidian isolates (Nosematidae: *Nosema*)from five important lepidopteran pests in Taiwan. J Invertebr Pathol. 2003; 83(1): 51–59.

15. García-Martínez J, Acinas SG, Antón AI, Rodríguez-Valera F. Use of the 16S-23S ribosomal genes spacer region in studies of prokaryotic diversity. J Microbiol Meth. 1999; 36(1-2): 55–64.

16. Couto I, Pereira S, Miragaia M, Sanches IS, de Lencastre H. Identification of clinical staphylococcal isolates from humans by internal transcribed spacer PCR. J Clin Microbiol. 2001; 39(9): 3099–03.

17. Tilsala-Timisjarvi A, Alatossava T. Characterization of the 16S-23S and 23S-5S rRNA intergenic spacer regions of dairy Propionibacteria and their identification with species-specific primers by PCR. Int J Food Microbiol. 2001; 68(1-2): 45–52.

18. Jeng RS, Svircev AM, Myers AL, Beliaeva L, Hunter DM, Hubbes M. The use of 16S and 16S-3S rDNA to easily detect and differentiate common Gram-negative Orchard Epiphytes. J Microbiol Meth. 2001; 44(1): 69–77.

19. Tan Z, Hurek T, Vinuesa P, Muller P, Ladha JK, Reinhold-Hurek B. Specific detection of *Bradyrhizobium* and *Rhizobium* strains colonizing rice (Oryza sativa)roots by 16S-23S ribosomal DNA intergenic spacer-targeted PCR. Appl Environ Microbiol.2001; 67(8): 3655–64.

20. Boyer SL, Flechtner VR, Johansen JR. Is the 16S-23S rRNA internal transcribed spacer region a good tool for use in molecular systematics and population genetics? A case study in cyanobacteria. Mol Biol Evol. 2001; 18(6):1057–69.

